# LRP1 and SORL1 regulate tau internalization and degradation and enhance tau seeding

**DOI:** 10.1101/2020.11.17.386581

**Authors:** Joanna M. Cooper, Aurelien Lathuiliere, Mary Migliorini, Allison L. Arai, Mashhood M. Wani, Simon Dujardin, Selen C. Muratoglu, Bradley T. Hyman, Dudley K. Strickland

## Abstract

The identification of the apoE receptor, LRP1, as an endocytic receptor for tau raises several questions about LRP1s’ role in tauopathies. Is internalized tau, like other LRP1 ligands, delivered to lysosomes for degradation? Does LRP1 internalize pathological tau leading to cytosolic seeding? Do other, related receptors participate in these processes? We confirm that LRP1 rapidly internalizes tau, leading to efficient lysosomal degradation. Employing brain homogenates from human Alzheimer brain, we find that LRP1 also mediates cytosolic tau seeding. We additionally found that another apoE receptor, *SORL1*, a gene implicated in AD risk, also mediates tau endocytosis, degradation, and release into the cytoplasm of seed competent species. These data suggest a role for these apoE receptors in tau uptake, as well as the competing processes of degradation and release to the cytoplasm. The balance of these processes may be fundamental to spread of neuropathology across the brain in Alzheimer disease.

## INTRODUCTION

In Alzheimer’s Disease (AD), neurofibrillary tangles (NFT) have a sequential accumulation pattern as the disease progresses that correlates with neuronal susceptibility and cognitive decline (Braak et al., 1991; Hyman et al., 1984; Serrano-Pozo et al., 2013). NFT consist of abnormal accumulations of excessively phosphorylated forms of the microtubule-associated protein tau within the cytoplasm of certain neurons. In mouse models of AD in which human mutant P301L tau is over-expressed in the entorhinal cortex, aggregated tau accumulates in brain regions with neuronal projections from the entorhinal cortex such as the dentate gyrus supporting the notion that the pathological tau protein can spread from one non-adjacent anatomical region of the brain to another (De Calignon et al., 2012; Harris et al., 2012; Liu et al., 2012; Polydoro et al., 2013). In this process, pathological forms of tau are thought to be transferred from cell to cell and “seed” aggregation of cytoplasmic tau by a prion-like templated misfolding of endogenous tau (DeVos et al., 2018; Kaufman et al., 2018; Medina et al., 2014; Takeda et al., 2015; Wegmann et al., 2019).

Mechanisms of tau spreading are not well understood, but the presence of extracellular tau in brain interstitial fluid (Yamada et al., 2011) led to the discovery that tau is constitutively secreted from neurons in a manner that is increased during neuronal activity and upon aging (Chai et al., 2012; Harrison et al., 2019; Huijbers et al., 2019; Merezhko et al., 2018; Pooler et al., 2013). Extracellular tau aggregates can transfer between co-cultured cells, are internalized by cells, and following endocytosis can induce fibrillization of intracellular tau (Frost et al., 2009; Swanson et al., 2017). Recent studies have provided evidence that the LDL-receptor protein 1 (LRP1) functions as an endocytic neuronal receptor for the uptake and spread of tau (Rauch et al., 2020).

LRP1 is a large endocytic and signaling receptor that binds numerous ligands and effectively delivers them to lysosomal compartments where they are degraded by lysosomal enzymes. LRP1 is highly expressed throughout the brain in neurons, astrocytes, microvascular endothelial cells, and microglia. It is a member of a large family of endocytic/signaling receptors, and is structurally similar also to sorting receptors such as SORL1, which is genetically linked to AD (Rogaeva et al., 2007). Both SORL1 and LRP1 are neuronal apolipoprotein E (apoE) receptors. ApoE genotype has a strong impact on development of late-onset AD, with the ε4 allele representing a risk factor and the ε2 allele being protective (Corder et al., 2008; Strittmatter et al., 1993).

The ability of LRP1 to mediate the endocytosis of tau raises questions of whether internalized tau, like other LRP1 ligands, is delivered to lysosomes for degradation. Further, it is not known if LRP1 participates in the processing of pathological forms of tau that lead to seeding. Moreover, it is unknown if LRP1 is the sole receptor for tau on the cell surface, or if other related molecules may also participate. The current study employed well-characterized cell lines deficient in LRP1 to address these questions. Our studies reveal that LRP1 efficiently delivers tau to lysosomal compartments for degradation, and that cells expressing LRP1, but not cells deficient in LRP1, also evidence endolysosomal escape of tau and promote tau seeding induced by soluble high molecular weight oligomeric tau fractions derived from brains of AD patients. In addition, we identified SORL1 as a second receptor that binds and internalizes tau and promotes tau seeding and find that the N1358S mutant of this receptor is deficient in these processes. These data put several molecules of clear importance in AD pathophysiology – LRP, apoE, SORL1, and tau – into a single molecular pathway involving uptake of pathological tau and seeding aggregation of intracytoplasmic tau.

## RESULTS

### LRP1 is an endocytic receptor for tau

Alternative splicing of the *MAPT* gene gives rise to six variants of tau protein, with the 2N4R variant being the largest. In our experiments the 2N4R variant was used unless otherwise noted. To test the hypothesis that LRP1 is responsible for mediating the cellular uptake of tau, we examined the endocytosis of ^125^I-labeled recombinant 2N4R tau in WT Chinese hamster ovary (CHO) cells and in CHO 13-5-1 cells, which are deficient in LRP1 (FitzGerald et al., 1995). The results (Fig 1a) reveal that the cellular uptake of ^125^I-labeled tau was significantly reduced, by about two thirds, in CHO cells lacking LRP1. The contribution of LRP1 to cellular-mediated uptake of tau was further confirmed by demonstrating that RAP, a high affinity LRP1 antagonist (Herz et al., 1991), prevented the uptake of tau in WT CHO cells. The time course of ^125^I-labeled tau surface binding and internalization in CHO WT and 13-5-1 cells reveals that both RAP and heparin, which has previously been reported to block tau internalization (Holmes et al., 2013), reduce the amount of ^125^I-labeled tau internalized in CHO WT cells, but neither had an effect on internalization of ^125^I-labeled tau in CHO 13-5-1 cells (Fig 1b). RAP and heparin also reduce ^125^I-labeled tau binding to the cell surface of WT CHO cells but have little effect on the binding of ^125^I-labeled tau to the surface of 13-5-1 cells (Fig 1b). The fact that CHO 13-5-1 cells appear to internalize small amounts of ^125^I-labeled tau that is not inhibited by either RAP or heparin confirms the existence of LRP1-independent pathways for tau internalization. In CHO cells, the alternative pathway(s) accounts for approximately 30% of tau internalization.

**Figure 1.**
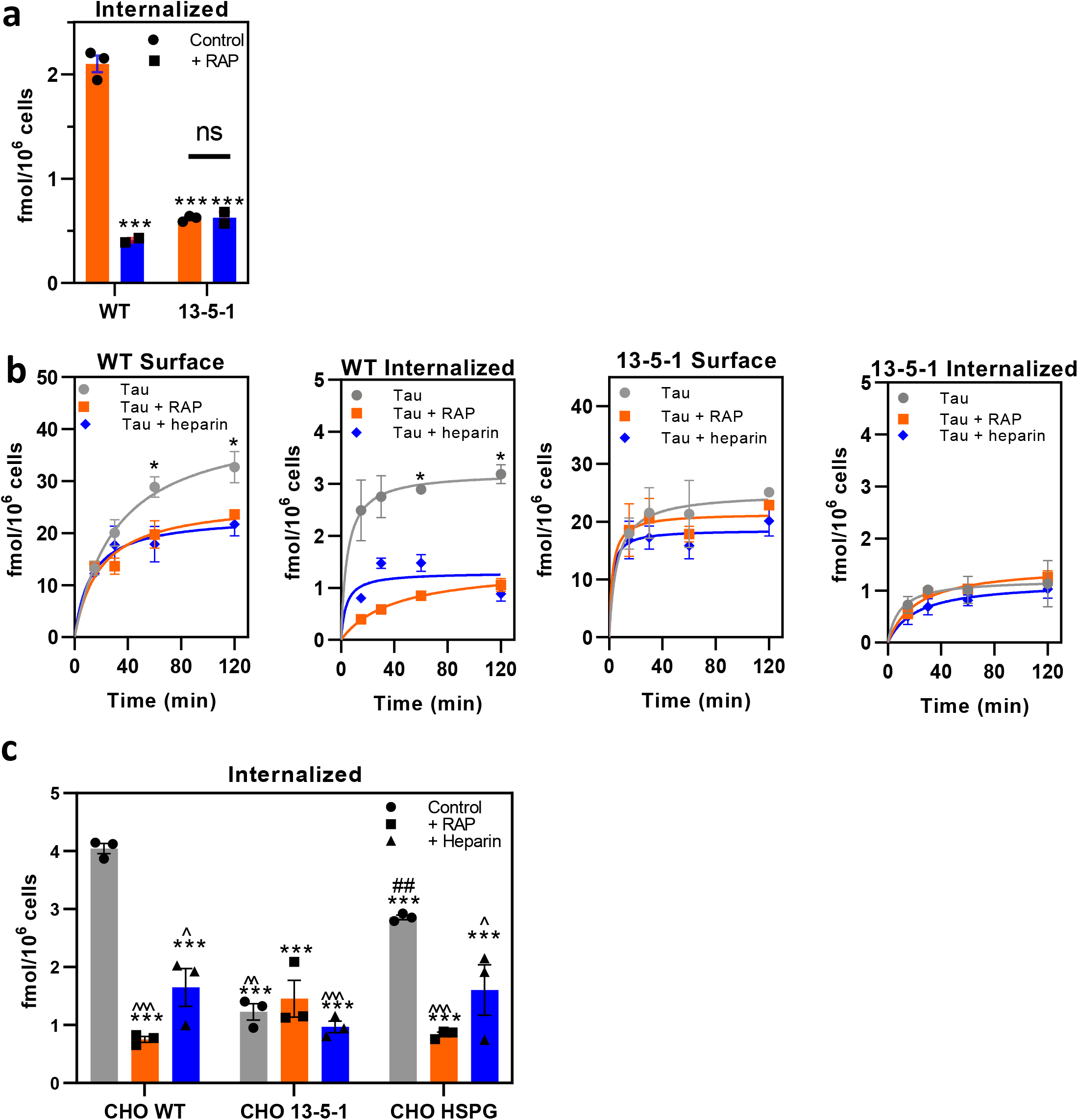
LRP1 is an endocytic receptor for tau. **a)** WT or LRP1-deficient 13-5-1, CHO cells were incubated with 20 nM ^125^I-labeled tau in the absence or presence of 1 μM RAP 2 h at 37°C and internalized measured. **(b)**. Time course for internalization of ^125^I-labled tau (20 nM) in CHO WT and CHO 13-5-1 cells in the presence or absence of RAP (1 μM) or heparin (20 μg/ml). Surface associated and internalized ^125^I-labeled tau were quantified; **(c)** WT, 13-5-1 and HSPG-deficient (CHO HSPG) CHO cells were incubated with 20 nM ^125^I-labeled tau in the absence or presence of RAP (1 μM) or heparin (20 μg/ml) at 37°C for 2 h and internalized measured. **(a,b,c)** Means ± SEM; two-way ANOVA followed by Sidak’s multiple comparisons test, **(a)**\*\*\**P*<0.0001 compared to WT control, *n*=3, (b) \**P*<0.0001 comparison of tau vs tau + RAP, *n*=3**(c)** significance reported compared to *CHO WT, #CHO 13-5-1, or ^CHO HSPG (1 symbol *P*<0.03; 2 symbols *P*<0.007; 3 symbols *P*<0.0001).

Previous studies have suggested that heparan sulfate proteoglycans (HSPG) regulate the cellular uptake of tau (Holmes et al., 2013; Rauch et al., 2018; Stopschinski et al., 2018). Thus, we also examined the uptake of ^125^I-labeled tau in CHO cells deficient in xylosyltransferase (Esko et al., 1985), an enzyme that catalyzes the first step in glycosaminoglycan synthesis. These cells lack HSPG as well as the glycosaminoglycans chondroitin sulfate and heparan sulfate. Our experiment compared the extent of ^125^I-labeled tau internalized these cells (labeled CHO HSPG) along with WT and LRP1-deficient CHO cells. The results of this experiment (Fig 1c) show a significant reduction in the amount ^125^I-labeled tau internalized in HSPG-deficient CHO cells when compared to WT CHO cells. RAP reduced the uptake of ^125^I-labelled tau in HSPG-deficient CHO cells, but as in our initial experiment has no significant effect on ^125^I-labeled tau uptake in LRP1-deficient CHO cells. These results suggest that glycosaminoglycans participate in the LRP1-mediated uptake of tau, similar to what we have observed for LRP1-mediated VLDL uptake induced by lipoprotein lipase (Chappell et al., 1994).

### LRP1-mediated endocytosis results in lysosomal degradation of tau

To determine if tau is degraded following internalization, we investigated the internalization and degradation of ^125^I-labeled tau in WI-38 fibroblasts, which express high levels of LRP1 and efficiently degrade other LRP1 ligands. The results of this experiment reveal that excess RAP reduces the extent of tau internalization and cellular-mediated degradation (Fig 2a). These experiments also revealed that internalized tau is effectively degraded in lysosomal compartments as demonstrated by the ability of the lysosomal inhibitor, chloroquine (CQ), to block its degradation. Curiously, chloroquine also reduced the amount of tau internalized (Fig 2a). Chloroquine inhibits endosomal acidification, and we hypothesize that this result is due to inefficient dissociation of tau from LRP1 within endosomal compartments resulting in recycling of the LRP1/tau complex.

**Figure 2.**
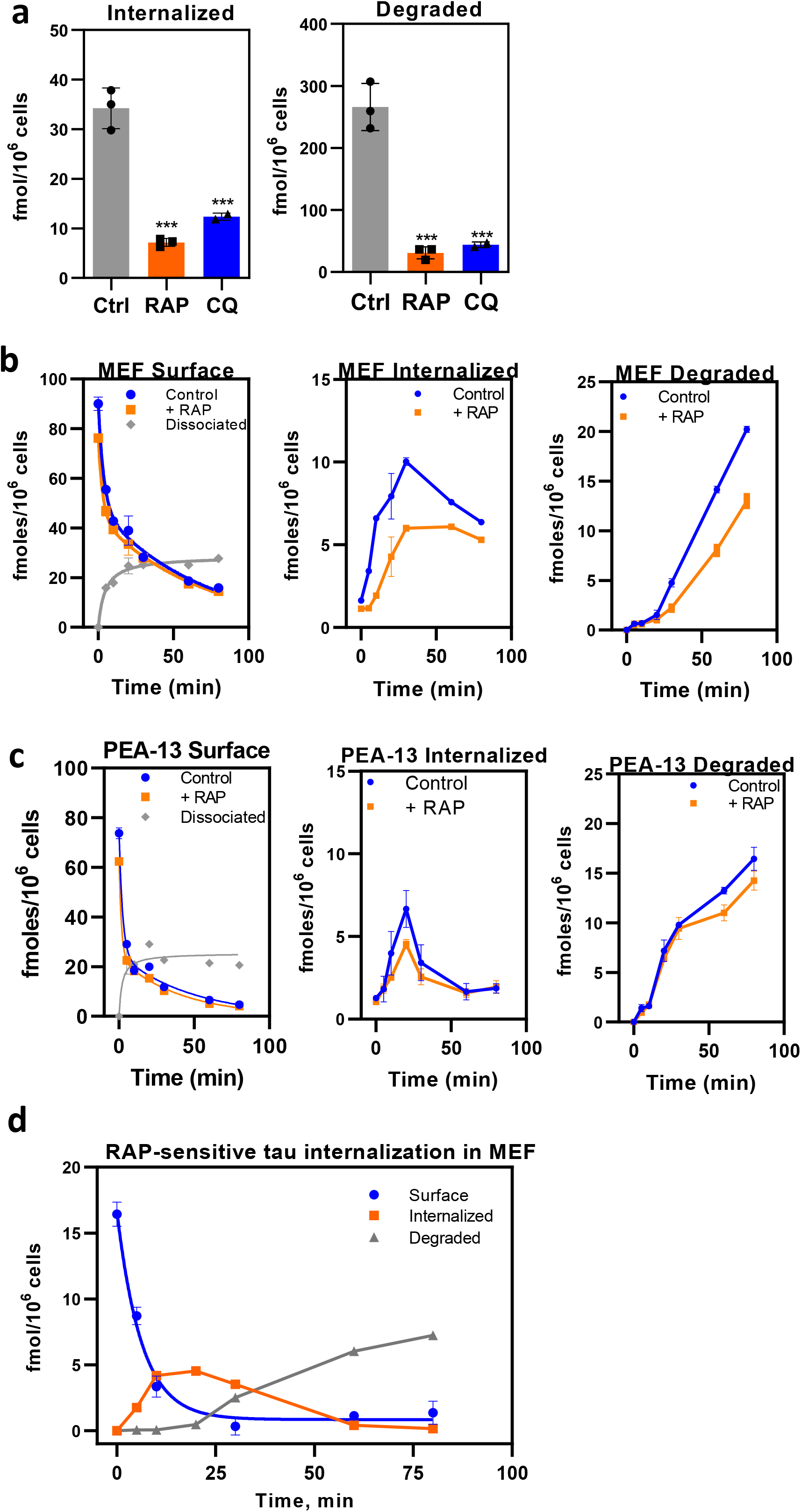
Single cycle endocytosis experiment reveals that tau is efficiently degraded following LRP1-mediated internalization. **a)** WI-38 cells were incubated with 20 nM ^125^I-labeled tau in the absence or presence of 1 μM RAP for 2 hours at 37°C, and the amount internalized (*right panel*) and degraded quantified (*left panel*). Degradation was measured in the presence or absence of or 100 μM chloroquine (CQ). (Means ± SEM; one-way ANOVA followed by Tukey’s multiple comparisons test (\*\*\**P*<0.001 compared to control, *n*=3). **(b,c)** MEF cells **(b)** or LRP1-deficient PEA-13 cells **(c)** were incubated with 20 nM ^125^I-labeled tau at 4°C for 2 hours in the presence or absence of 1 μM RAP, then the media was replaced with warm assay media ± RAP, and cells were incubated at 37°C for specified times. The amounts of surface bound, internalized, degraded, and dissociated ^125^I-labeled tau were quantified. **(d)** RAP-sensitive surface, internalized and degraded ^125^I-labeled tau in MEF cells was calculated from the data in b by subtracting the RAP inhibitable tau uptake from the total.

To examine the cellular processing of tau mediated by LRP1, we performed a single cycle endocytosis experiment. In this experiment, ^125^I-labeled tau was incubated with mouse embryonic fibroblasts expressing LRP1 (MEF) (Fig 2b) or lacking LRP1 (PEA-13) (Fig 2c), at 4°C for 1 hr in the absence or presence of RAP to block LRP1-mediated uptake. After washing, the media was replaced with fresh media in the absence or presence of RAP and maintained at 37°C to trigger endocytosis, and the cell-associated, internalized and degraded tau was quantified. In LRP1 expressing cells, ^125^I-labeled tau disappears rapidly from the cell surface, with some of it dissociating into the media (Fig 2b, *left panel*). A significant portion of the ^125^I-labeled tau is internalized and with time is degraded (Fig 2b, *middle and right panels*). Some of the surface binding, internalization and degradation in these cells is blocked by RAP. A similar amount of binding to the cell surface was observed in LRP1-deficient (PEA-13) cells, although much less was internalized and degraded, and none was RAP sensitive (Fig 2c). The presence of some residual uptake and degradation confirms, in a second cell system, the existence of LRP1-independent mechanism(s) for tau internalization. To quantify the surface binding, internalization and degradation mediated by LRP1, RAP sensitive surface, internalized and degraded ^125^I-labeled tau was calculated from the data by subtracting the RAP-sensitive component from the data of Fig 2b. The results reveal a rapid internalization of tau with a half-life of 4 min. After a lag period of approximately 15 min, tau degradation was detected (Fig 2d). These data reveal that LRP1-mediated endocytosis results in effective trafficking of tau to lysosomal compartments and subsequent degradation in these cells.

### LRP1 also mediates tau internalization and degradation in immortalized central nervous system derived cell cultures

To determine if tau internalization and degradation also occurs in brain relevant cell cultures, we also examined the internalization and degradation of tau in H4 neuroglioma cells. After 2h of incubation, the results reveal that these cells also mediate the internalization and degradation of ^125^I-labeled tau in a RAP sensitive manner (Supplementary Fig 1a *left panel*). Since RAP potentially interacts with several members of the LDL receptor family, the role of LRP1 in this process is revealed by the ability of anti-LRP1 IgG (R2629) to reduce the amount of tau internalized (Supplementary Fig 1a, *left panel*). To detect degraded tau, we extended the incubation period to 24 h, and confirmed that a significant amount of ^125^I-labeled tau is internalized and degraded in a RAP-sensitive manner (Supplementary Fig 1a, *middle and right panel*). We next examined the internalization of tau in the neuroblastoma cell line SH-SY5Y, to investigate tau uptake in a second neuronal cell culture, by employing immunofluorescence with the goal of determining if tau co-localizes with LRP1 during endocytosis. In these experiments, functional LRP1 was labeled with a monoclonal antibody that recognizes the LRP1 light chain and does not dissociate from the receptor during endosomal trafficking and receptor recycling (Muratoglu et al., 2010). We then incubated the live cells with fluorescently labeled tau. The results demonstrate co-localization of LRP1 and tau within endosomal compartments (Supplementary Fig 1b).

### Tau binding to purified LRP1 occurs with high affinity but is not dissociated by low pH in vitro

To extend our cell-based results, we investigated the binding of tau to purified LRP1. Our initial experiments utilized an ELISA in which we immobilized purified LRP1 on the surface of microtiter plates and measured the ability of increasing concentrations of tau to bind to the LRP1 coated wells. As controls, we also measured the binding of tau to LRP1 in the presence of RAP and to BSA-coated wells. The results of this experiment are shown in Fig 3a and confirm RAP-inhibitable binding of tau to LRP1. Further, the results reveal that tau selectively binds to LRP1-coated wells, but not to BSA-coated wells.

**Figure 3.**
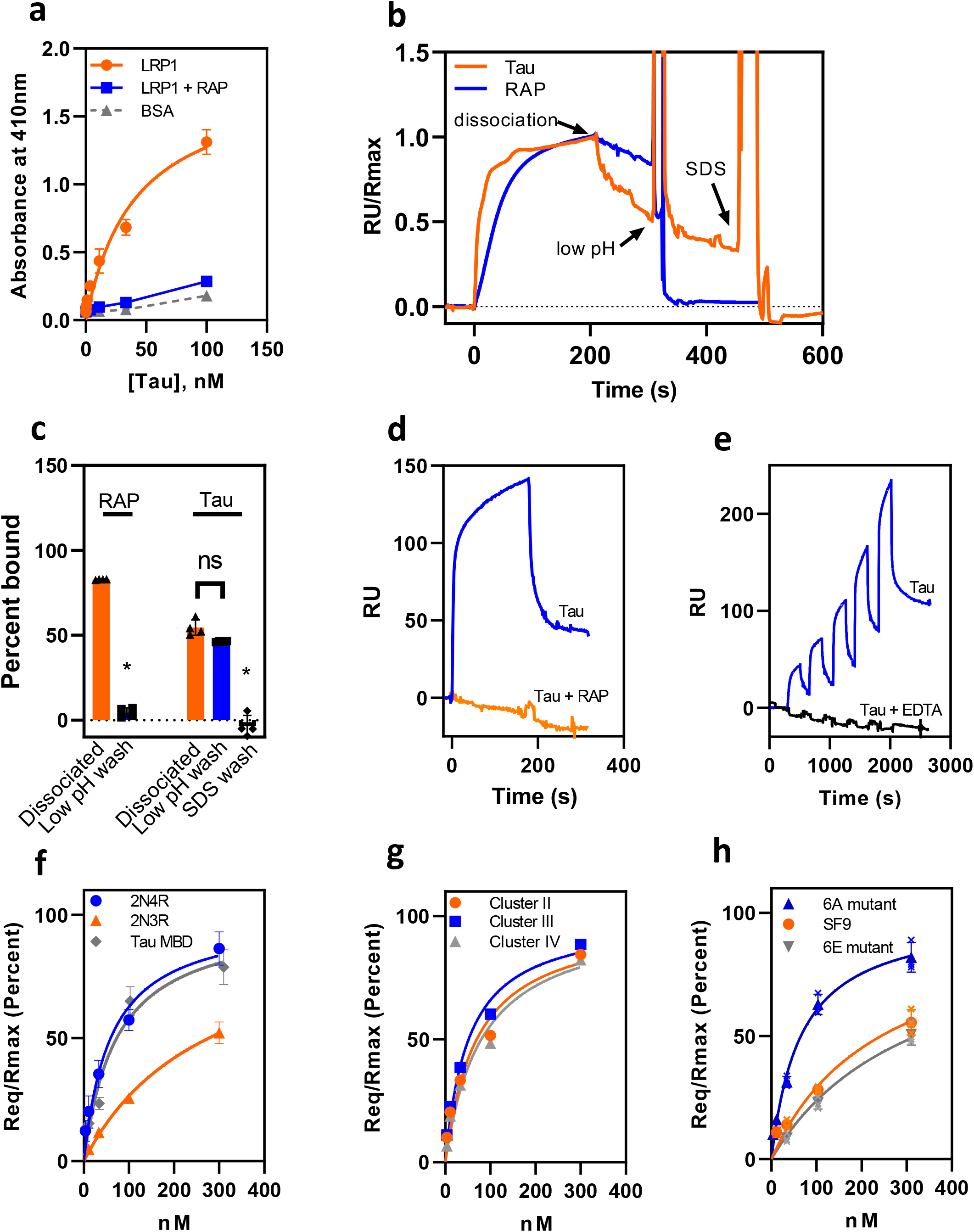
SPR analysis confirms high affinity binding of tau to LRP1. **(a)** ELISA measuring binding of 2N4R tau to LRP1 in the presence or absence of RAP (circles) or BSA (triangles) Shown are means ± SEM, n=3. (**b)** Tau binding is not sensitive to pH. Tau (500 nM) or RAP (20 nM) was allowed to bind to LRP1 coupled to a SPR chip followed by dissociation. The chip was then regenerated with low pH followed by 0.5% SDS. **(c)** Percent of 20 nM RAP and 500 nM tau remaining bound to full-length LRP1 was calculated following 100 sec dissociation, low pH wash SDS wash. Shown are mean ± SEM of three independent replicate experiments. (RAP binding: t-test, \**P*<0.0001, *n*=3,. Tau binding: one-way ANOVA followed by Tukey multiple comparisons test, \**P*<0.0001, *n*=3. (d) Inhibition of tau binding by RAP as assessed by co-injection experiment. **(e)** Single cycle kinetic experiment quantifying binding of monomeric tau (3.8, 11.5, 34.4, 103.3, 310 nM) to LRP1 in the presence of Ca^2+^ or EDTA. **(f)** Binding of tau isoforms 2N4R, 2N3R, and tau MBD to LRP1 assessed by SPR equilibrium analysis **(g)** Binding of monomeric tau to LRP1 clusters II, III, or IV by SPR equilibrium analysis **(h)** The binding of tau produced by Sf9 cells along with two mutant forms of tau to full-length human LRP1 were measured by SPR; 6A, (T181, S199, S202, S396, S400 and S404 are all converted to alanine), 6E, in which all of these residues are converted to glutamic acid.

To quantify the interaction of tau with LRP1 we utilized surface plasmon resonance (SPR) experiments. In these experiments, we observed high affinity tau binding to LRP1-coupled chips. Interestingly, we found that tau remained bound to LRP1 during low-pH mediated regeneration of our LRP1-coupled SPR chip, and was only released upon washing with SDS solutions (Fig 3b,c). This is not the case for other LRP1 ligands, such as RAP, which readily dissociates in the presence of a low pH wash with 100 mM phosphoric acid at pH ~2.5, which is routinely used to regenerate LRP1-coated SPR chips after testing binding properties with other LRP1 ligands (Fig 3b,c). These data reveal that binding of tau to LRP1 is not as sensitive to low pH as other LRP1 ligands, and thus may not fully dissociate from LRP1 in endosomal compartments. To confirm that RAP blocks the binding of tau to the LRP1-coated SPR surface, we conducted competition experiments in which we quantified the amount of tau binding in the presence of excess RAP. The results confirm that RAP effectively competes for the binding of tau to the LRP1-coated SPR surface (Fig 3d). To determine the KD for the interaction of tau with LRP1, we used single cycle kinetic measurements which do not require surface regeneration (Fig 3e). We confirmed the specificity of the interaction by demonstrating that the binding of tau to the LRP1-coated chip was ablated in the presence of EDTA, which chelates the essential Ca^2+^ ions necessary to stabilize the LDLa ligand binding repeats which are critical for ligand binding by this class of receptors (Fig 3e, *black lines)*. To determine the KD of this interaction, we fit the individual data to a pseudo–first-order process to obtain values of Req for each concentration of tau isoforms (2N4R, 2N3R and tau microtubule-binding domain), and then plotted the Req values as function of total concentration of tau (Fig 3f). Nonlinear regression analysis of the plot revealed a KD value of 60 ± 8 nM for the 2N4R tau isoform, a value comparable to other LRP1 ligands such as soluble forms of APP(Kounnas, Moir, et al., 1995) or hepatic lipase(Kounnas, Chappell, et al., 1995).

Tau contains two major domains: an N-terminal “projection” domain containing the alternatively spliced N1 and N2 regions and the C-terminal microtubule binding domain containing four highly conserved repeat regions, R1-R4, which binds to microtubules (Nizynski et al., 2017). Interestingly, the 2N3R isoform of tau which lacks the second microtubule binding repeat (R2) encoded by the alternatively spliced exon 10, bound to LRP1 with considerably weaker affinity (KD = 278 ± 55 nM, Fig 3f) suggesting that the R2 domain of tau contributes to the interaction of tau with LRP1. We also quantified the interaction of the microtubule binding domain (R1-R4, leu243-glu372) with LRP1 using SPR measurements and the results of these experiments reveal that this region of tau binds to LRP1 with an affinity similar to the intact molecule (KD = 73 ± 18 nM) (Fig 3f). These results suggest that the binding region of tau that interacts with LRP1 is localized to the R2 domain of tau and indicates that the microtubule binding domain alone is sufficient for binding to LRP1.

The ligand binding regions of LRP1 are mainly localized to clusters of LDLa repeats, termed clusters I, II, III and IV. To determine which region of LRP1 is involved in tau binding, we investigated the binding of tau to clusters II, III and IV immobilized on SPR chips. The results of a single cycle kinetic experiment confirm that tau readily binds to clusters II, III and IV with KD values of 69 ± 25, 52 ± 14 and 81 ± 29 nM, respectively (Fig 3g). The binding of tau to all three clusters of LRP1 with similar affinity is unusual as most ligands prefer to bind to clusters II or IV (Neels et al., 1999). The lack of tau binding preference amongst all the LRP1 clusters may indicate some cooperativity in the binding of tau to cellular LRP1 that is not detectable on LRP1 covalently crosslinked to an SPR surface.

### Phosphorylation of tau reduces its affinity for LRP1

Tau contains multiple serine, threonine and tyrosine phosphorylation sites that have been extensively studied as phosphorylation is a common post-translational modification of tau (Bramblett et al., 1993; Hanger et al., 2007; Mandelkow et al., 1995). Phosphorylated forms of tau are detected in tau aggregates in Alzheimer disease and other tauopathies, as well as in the high molecular weight soluble forms of tau that have been implicated in neuronal uptake and propagation of seed competent species (Grundke-Iqbal et al., 1986; Ihara et al., 1986; Iqbal et al., 1989; Takeda et al., 2015). In addition, tau phosphorylation at specific residues regulates tau’s function, regulates its subcellular localization (Pooler et al., 2012; Sultan et al., 2011; Tang et al., 2015) and reduces its affinity for microtubules (Biernat et al., 1993; Jameson et al., 1980). Therefore, we tested the hypothesis that phosphorylation of tau might alter its binding to LRP1 by examining the binding of recombinant tau produced by Sf9 insect cells which produce well characterized hyperphosphorylated forms of tau (Mair et al., 2016; Tepper et al., 2014). We found that hyperphosphorylated tau produced by Sf9 cells bound LRP1 with a 4-fold weaker affinity (KD = 243 ± 17 nM) (Fig 3h). We also examined the binding of two recombinant (E. Coli produced) mutant forms of tau to LRP1: mutant 6A, in which T181, S199, S202, S396, S400 and S404 are all converted to alanine, and mutant, 6E, in which all of these residues are converted to the phosphomimetic glutamic acid. These specific residues have been found to be phosphorylated in both normal and AD brains (Šimić et al., 2016). Our results reveal that while the 6A mutant binds to LRP1 with a KD value similar to that of WT tau (KD = 65 ± 4 nM), the 6E mutant binds to LRP1 with 5-fold weaker affinity (KD = 321 ± 17 nM) (Fig 3h). Together, the results of our studies reveal that phosphorylated forms of tau bind to LRP1 with significantly lower affinity. Since phosphorylation of tau is generally associated with increased tau pathology, reduced binding of phosphorylated tau to LRP1 suggests that the LRP1-mediated pathway is less tightly bound to LRP1 compared to presumably less modified wild type tau.

### LRP1 provides a mechanism of uptake that supports tau proteopathic seeding in the cytoplasm

LRP1 efficiently mediates tau uptake and degradation using recombinant forms of the protein, and certain post-translational modifications appear to diminish tau-LRP1 interactions. Key to the biological import of these observations, then, is the question of whether the robustly modified tau present in human AD brain is also an LRP1 ligand. Moreover, key to understanding LRP1’s potential role in tau propagation across cells is whether LRP1 both mediates uptake of human brain derived tau, and also allows it to escape from endosomal/lysosomal systems, providing misfolded tau access to the cytoplasm to lead to seeding of endogenous tau. To determine if the LRP1-mediated uptake of pathological forms of tau results in tau seeding, we conducted experiments in which brain samples isolated from a Braak VI AD patient and a healthy control were incubated with CHO WT or CHO 13-5-1 cells that had been transfected with a tau seeding bioreporter construct. To detect tau seeding, we used a FRET-based biosensor assay analogous to that described by Holmes et al. (Holmes et al., 2014) by transfecting both cells lines with a pcDNA3 plasmid containing a construct that encoded residues 344-378 of human P301L mutant tau fused to either mTurquoise2 or to Neon Green. The results of this experiment reveal that incubation of CHO WT cells with brain extract from an AD patient induces tau seeding as revealed by increased FRET, whereas incubation of brain extract from a healthy control has little effect on tau seeding (Fig 4a). In contrast, incubation of brain homogenate from AD patients with LRP1-deficient CHO 13-5-1 cells results in only marginally detectable amount of tau seeding. As a control for these experiments, when the plasma membrane of either cell line was permeabilized with lipofectamine, tau seeding occurred (Fig 4b) to an equivalent extent. CHO WT cells also demonstrate tau seeding induced by HMW SEC tau fractions of brain extract from AD patients while tau seeding is significantly reduced, though still measurable, in LRP1-deficient CHO 13-5-1 cells (Fig 4c). In WT CHO cells, the extent of tau seeding resulting from incubation with HMW SEC tau fractions of brain extracts from AD patients is reduced in the presence of RAP and anti-LRP1 antibodies (Fig 4d). Together, these experiments reveal that LRP1 supports the uptake and endolysosomal escape of pathological forms of tau resulting in tau seeding. We also noted that in CHO WT cells, incubation with chloroquine significantly increased tau seeding when the cells were incubated with HMW SEC fractions from brain extracts of AD patients (Fig 4e). This suggests that preventing tau degradation in the lysosome enhances the likelihood that seed competent tau escapes to the cytoplasm where it can interact with the biosensor molecule.

**Figure 4.**
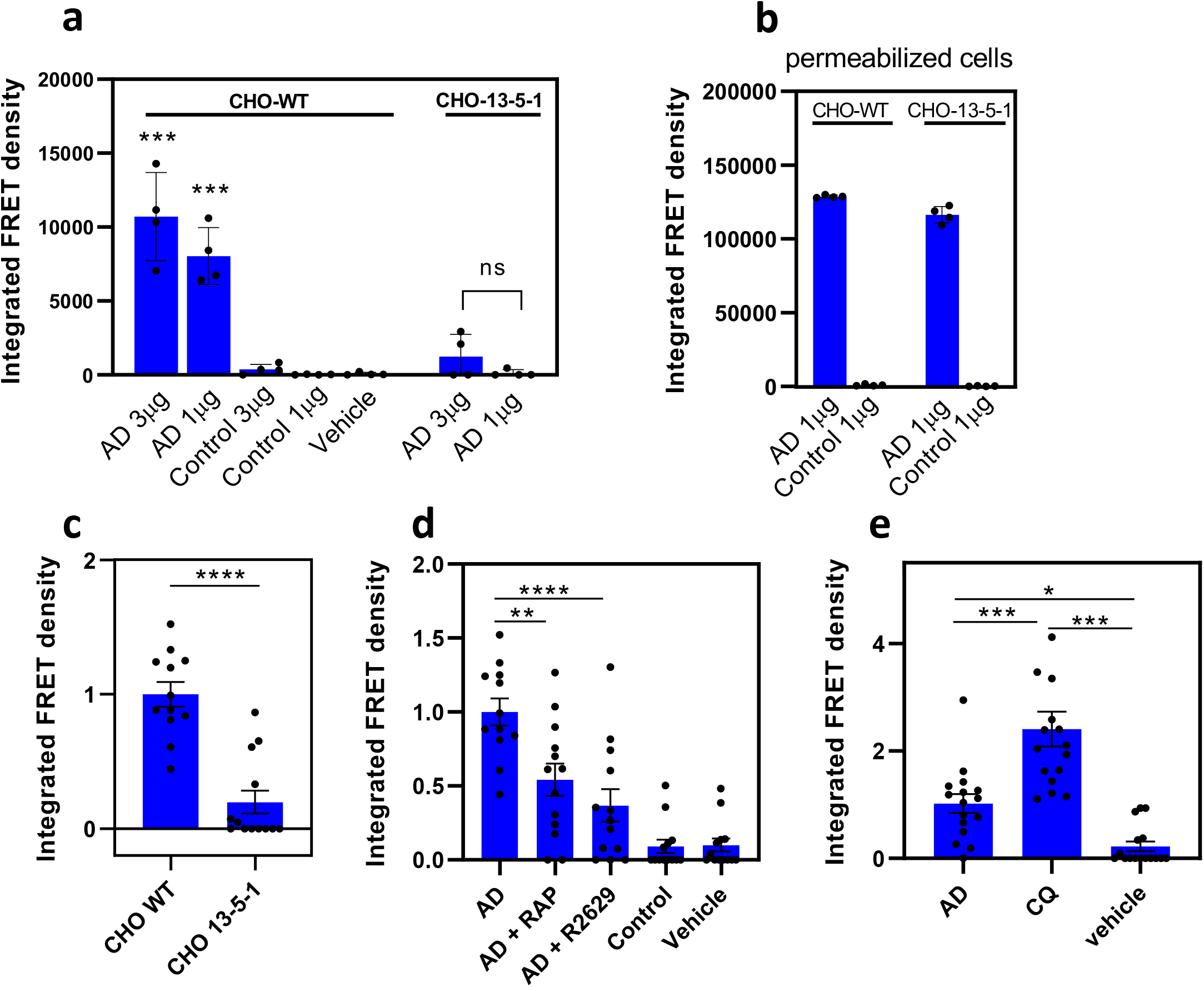
LRP1 mediates tau seeding. CHO WT and 13-5-1 cells were transfected with pcDNA3 plasmid containing a construct that encoded residues 344-378 of human P301L mutant tau fused to either mTurquoise2 or to Neon Green and **(a)** incubated with 1-3 μg of human brain homogenate from an Alzheimer’s patient or from a healthy control for 24 hours. **(b)** As a positive control, 1% lipofectamine 2000 was added to the wells. Tau seeding was quantified by multiplying the percent of FRET-positive cells by the median florescence intensity of those cells. Each condition was performed in at least quadruplicate and data was analyzed using FlowJo software. **(a & b)** Means ± SD, one-way ANOVA followed by Tukey multiple comparisons test \*\*\**P*<0.0001 compared to vehicle control. **(c)** Transfected CHO WT and 13-5-1 cells were incubated with HMW SEC fractions from human brain of an AD patient, shown are means ± SD, t-test *n*=13, \*\*\*\**P*<0.0001. CHO WT cells transfected with the tau FRET reporter system were **(d)** incubated with HMW SEC fractions from human brain of an AD patient in the presence or absence of 1 μM RAP or LRP1 antibodies (R2629), **(e)** incubated with HMW SEC fractions from human brain of an AD patient with 100 μM chloroquine (CQ) (Means ± SD, one-way ANOVA followed by Tukey’s multiple comparisons test \*\*\**P*<0.0001, \**P*<0.03.

### SORL1 also binds and mediates the endocytosis of tau

Our studies have confirmed the existence of multiple pathways for mediating the endocytosis of tau, and we next initiated studies to identify additional receptors involved in this process. The studies of Rauch et al. (Rauch et al., 2020) suggested that LRP1B, LRP2, LRP5, LRP8, LDLR, and VLDLR are not involved in tau uptake. Since we previously found that SORL1 associated with LRP1 (Spoelgen et al., 2010), and since SORL1 is genetically associated with AD, we examined the potential of this receptor to mediate tau uptake. CHO-WT and CHO 13-5-1 cells were transfected with a plasmid containing a construct that encoded *SORL1*, using cells incubated with transfection reagent only (“mock”). Transfection efficiency was validated via western blot (Supplementary Figure 2a). The results reveal that both CHO WT (Fig 5a) and CHO 13-5-1 cells (Fig 5b) expressing SORL1 bind more ^125^I-labeled tau on the cell surface and show a dramatic increase in the amount of tau internalized in a process inhibited by RAP. Direct binding of tau to recombinant VS10P domain of tau (residues 82-753) was confirmed by SPR measurements, which revealed a KD value of 41 ± 9 nM (Fig 5c).

**Figure 5.**
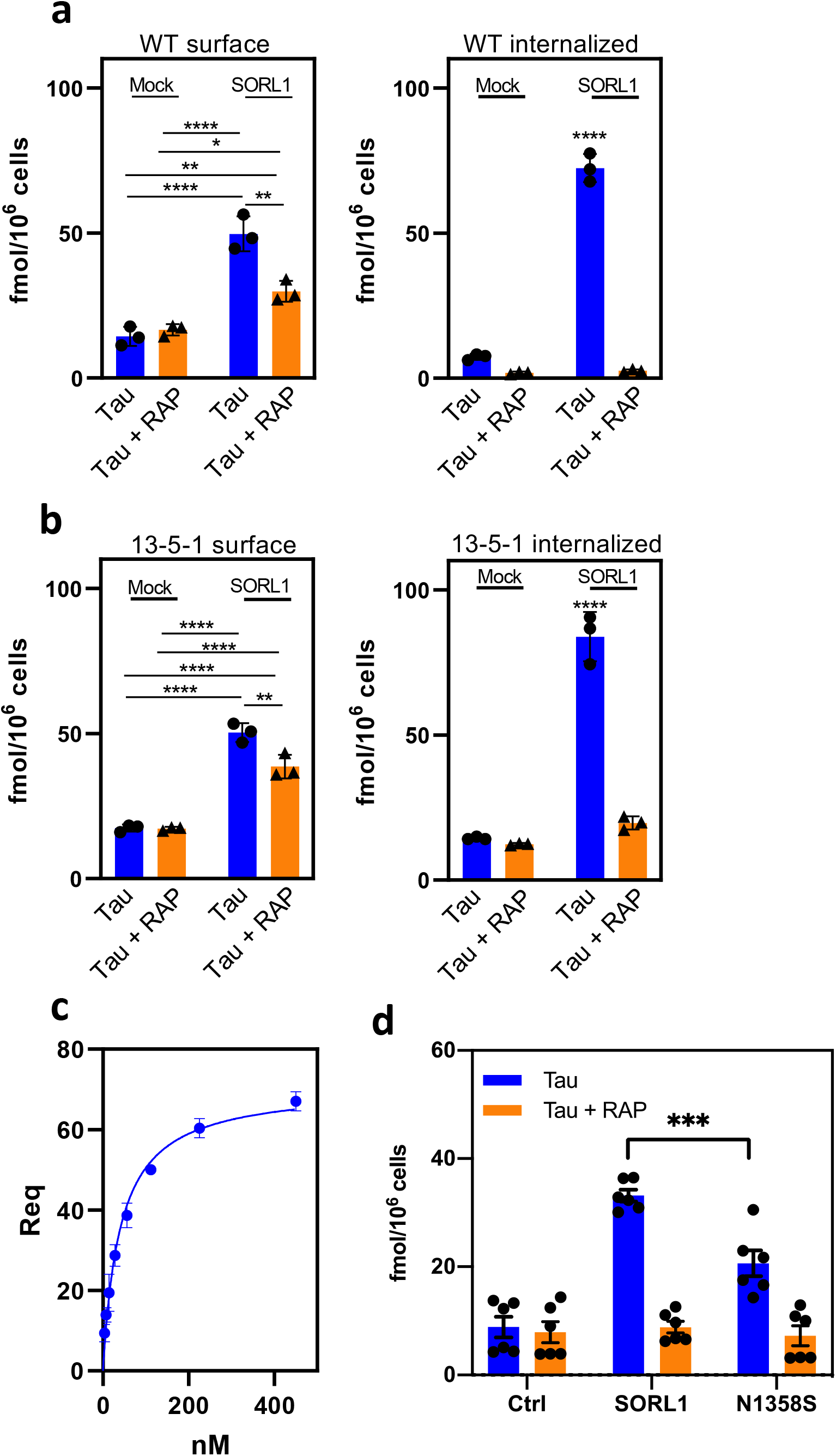
SORL1 mediates tau internalization. CHO WT **(a)** or CHO 13-5-1 **(b)** cells were transfected with *SORL1* plasmid, incubated with 20 nM ^125^I-labeled tau in the presence or absence of 1 μM RAP for 2 hours, and then surface bound and internalized tau was quantified. **(c)** binding of increasing concentrations of 2N4R tau to SORL1 VSP10 domain coupled to a Biacore CM5 sensor chip. **(d)** Internalized tau in CHO 13-5-1 cells transfected with *SORL1* or N1358S mutated *SORL1* plasmid, incubated with 20 nM ^125^I-labeled tau in the presence or absence of 1 μM RAP for 2 hours (*n*=6). (Means ± SD, two-way ANOVA followed by Tukey multiple comparisons test \*\*\*\**P*<0.0001, \*\**P*<0.001, \**P*<0.01, *n*=3).

In a separate experiment we transfected CHO 13-5-1 cells with a plasmid expressing SORL1 with the N1358S mutation, a SORL1 variant identified by exome sequencing in patients with early onset AD, located in the complement repeat 7 of SORL1 (Pottier et al., 2012). CHO 13-5-1 cells transfected with plasmid containing SORL1 harboring the N1358S mutation internalized less tau than those transfected with plasmid containing WT SORL1 (Fig 5d). Transfection efficiency was validated via western blotting.

### LRP1 and SORL1 increase tau seeding in HEK293T cells

HEK293T cells that stably express the P301S FRET biosensor are commonly used to assay tau seeding activity. Initially, we investigated the functional levels of LRP1 in HEK293T cells by comparing uptake of ^125^I-labeled alpha-2-macroglobulin, a well-characterized LRP1 ligand, in CHO WT, LRP1-deficient CHO 13-5-1, and HEK293T cells. The results of this experiment reveal very little alpha-2-macroglobulin uptake in HEK293T cells consistent with Western blots showing only very low levels of LRP1 in these cells (Supplemental Fig 2b). As expected, no alpha-2-macroglobulin internalization occurs in the LRP1-deficient CHO 13-5-1 (Fig 6a). HEK293T cells were transfected with human LRP1 and the amount of ^125^I-labeled tau uptake quantified. Transfection efficiency was validated via western blot (Supplementary Fig 2c). The results of this experiment demonstrate that the transfected cells readily internalize increased levels of tau in a process inhibited by RAP (Fig 6a, *left panel*).

**Figure 6.**
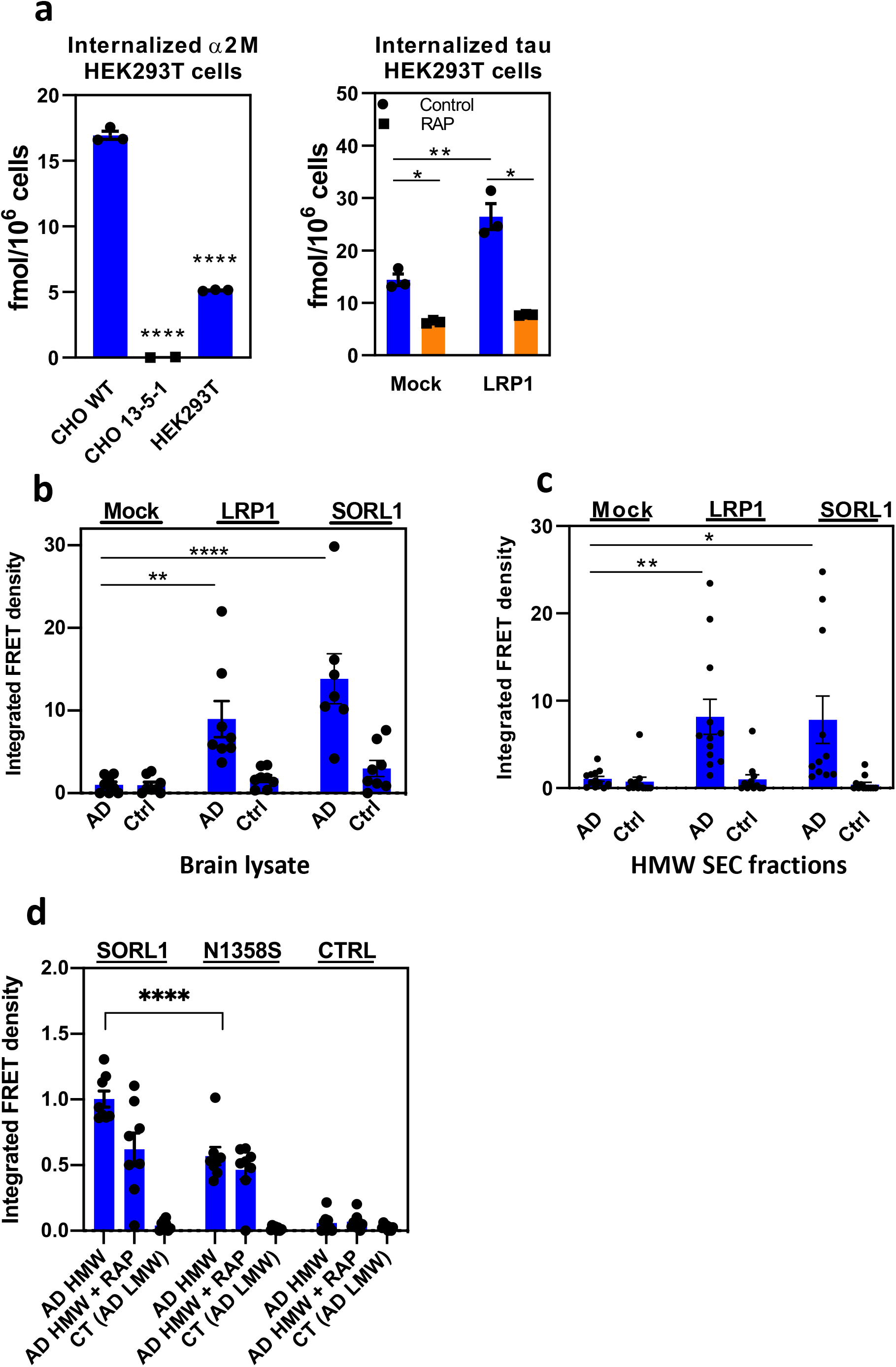
LRP1 and SORL1 mediate tau internalization and seeding in HEK293T cells. **(a)***Left panel*: CHO WT, 13-5-1, and HEK293T cells were incubated with 5 nM ^125^I-labelled alpha-2-macroglobulin for 2 hours and internalized alpha-2-macroglobulin was quantified. *Right panel:* HEK293T cells were transfected with LRP1 or mock transfected. 24 hours post transfection cells were incubated with 20nM tau in the presence or absence of 1 μM RAP for 2 hours, and internalized tau was quantified (*n*=3). HEK293T FRET reporter cells were transfected with *LRP1* or *SORL1*, then incubated with **(b)** human brain homogenate from an Alzheimer’s patient (n=8) or (c) HMW SEC fractions from AD patient brain (*n*=12). **(d)** HEK293T FRET reporter cells were transfected with *SORL1* or N1358S *SORL1*, then incubated with HMW SEC fractions from AD patient brain (*n=*8). (Means ± SEM, one way ANOVA (a) or two way ANOVA (b-d) followed by Tukey’s multiple comparisons test. * *P*<0.05, \*\**P*<0.01, \*\*\*\**P*<0.0001)

In the final series of experiments we utilized HEK293T biosensor cells(Holmes et al., 2014) to determine if LRP1 and SORL1 can mediate tau seeding in these cells. These cells were transfected with either LRP1 or SORL1 and incubated with brain lysate from AD patients (Fig 6b) or with HMW SEC fractions isolated from brains of AD patients (Fig 6c). The results of these experiments reveal that increased expression of LRP1 or SORL1 promotes a significant increase in tau seeding induced by brain lysates or HMW SEC fractions from AD brains. Cells transfected with N1358S SORL1 showed reduced tau seeding as compared to those transfected with WT SORL1 when incubated with HMW SEC fractions from AD brains.

## DISCUSSION

LRP1 and SORL1 are endocytic receptors that have been implicated in trafficking and in endosomal-lysosomal degradation. They recognizes numerous ligands and mediate their internalization and delivery to lysosomal compartments for processing and degradation. The overall objective of the current investigation was to quantify the extent to which tau is delivered to lysosomal compartments upon association with LRP1 and to determine if LRP1 can promote tau seeding, an event that is believed to occur only if seed competent tau escapes degradation in the lysosome and accesses the cytoplasm. Using cell lines deficient in LRP1 or antagonists to block LRP1 function, we quantified the uptake and cellular processing of 125I-labeled tau in multiple cell lines. Our results reveal that cells deficient in LRP1 are not effective in internalizing tau, although some residual tau uptake could be detected even in LRP deficient cells. As noted below, some of this residual uptake may be due to SORL1. By employing single cycle endocytosis experiments, our results also reveal that LRP1 efficiently delivers tau to lysosomes where the ligand is degraded.

Our cell-based data are supported by SPR experiments confirming high affinity binding of monomeric forms of tau to LRP1. Furthermore, our data confirm that the microtubule binding domain of tau binds tightly to LRP1, and reveal that ligand binding clusters II, III, and IV of LRP1 are capable of binding tau. It is somewhat unusual for an LRP1 ligand to be recognized by all three LRP1 ligand binding repeats and suggests that tau may bind to dimeric forms of LRP1 on cells resulting in increased affinity via avidity effects. Our SPR experiments also noted that unlike other LRP1 ligands we have investigated, the dissociation of monomeric forms of tau from LRP1 appears to be insensitive to pH, suggesting that tau may not dissociate from LRP1 within the low pH environment of endosomal compartments. The consequence of the failure of ligands to dissociate from LRP1 within endosomal compartments is not known at this time. In the case of the LDL receptor, failure of the ligand proprotein convertase subtilisin/kexin type 9 (PCSK9) to dissociate from the LDL receptor in endosomes results in altered receptor trafficking that leads to lysosomal-mediated receptor degradation (Leren, 2014).

Our results support the recent studies of Rauch et al. (Rauch et al., 2020) and Evans et al (Evans et al., 2020) who demonstrated that LRP1 can mediate the internalization of tau using flow cytometry approaches. In addition, the Rauch et al. (Rauch et al., 2020) study employed a unique adeno-associated virus construct (expressing GFP-2A-hTau) that is capable of discriminating between transduced cells and cells internalizing secreted tau (Wegmann et al., 2019) to demonstrate that neuronal LRP1 expressed in the brain of mice readily internalizes neuronally secreted tau, but this assay does not distinguish among the various post-translationally modified forms of tau that are present in the CNS, and does not distinguish between tau uptake and tau seeding. To determine if LRP1 can promote tau seeding, we incubated LRP1-expressing and LRP1-deficient cells with human brain homogenates from AD and from healthy controls and used a FRET-based biosensor assay to detect seeding. These studies revealed that LRP1-expressing cells promoted tau seeding when incubated with homogenates from AD patients but not homogenates from healthy controls. In addition, expression of LRP1 in HEK293T cells, which have low levels of LRP1, results in tau seeding when incubated with human brain homogenates from AD patients. Together, these results confirm that LRP1 is sufficient to promote tau seeding. While the LRP1-mediated tau uptake results in effective delivery of tau to lysosomal compartments resulting in degradation, future studies are required to determine where and how pathogenic forms of tau are processed. It is conceivable that tau may escape from the endosomal pathway. As a precedent for this, the LRP1 ligand, *Pseudomonas* exotoxin A, is cleaved within endosomal compartments releasing a 37 kDa domain that is translocated to the cytosol where it inhibits ADP ribosylation of elongation factor 2 (Kounnas et al., 1992). Interestingly, arresting endo-lysosomal trafficking with chloroquine increases the escape of tau seeds to the cytosol, possibly by extending the residence time in a susceptible exit compartment. It also remains unknown whether co-receptors exist that facilitate the uptake and endo-lysosomal escape of multimeric or other post-translationally modified forms of tau.

Our data highlight that the interaction of tau and LRP1 may be influenced by post translational modifications in tau since phosphorylated forms of tau bound much more weakly to LRP1 than unphosphorylated tau forms. The weaker affinity of these pathologic forms of tau for LRP1 may have important consequences on trafficking of tau and may allow modified tau to dissociate from LRP1 within early endosomes to facilitate endosomal escape. It is interesting to highlight in this regard that experiments employing hybrid constructs of *Pseudomonas* exotoxin A noted an inverse correlation between LRP1 binding affinity and toxicity (Zdanovsky et al., 1996), which requires endosomal escape of the toxin. For example, replacement of the receptor binding domain on the toxin with RAP generated a hybrid toxin with substantial increase in affinity for LRP1 but was much less toxic to cells. Presumably, the higher affinity of the hybrid toxin for LRP1 resulted in more efficient delivery to lysosomal compartments where the toxin was degraded.

Early studies recognized the importance of basic amino acids on the ligand that seemed critical for LRP1 receptor binding (Arandjelovic et al., 2005; Mahley et al., 1977; M. M. Migliorini et al., 2003; Prasad et al., 2016; Weisgraber et al., 1978). When the structure of two complement-like repeats (CR) from the LDL receptor in complex with the third domain of RAP (Fisher et al., 2006) was solved, a canonical model for ligand binding to LRP1 was revealed in which acidic residues from each CR formed an acidic pocket for in which two lysine side chains (K256 and K270) from RAP are docked. The acidic pocket on the receptors is stabilized by a calcium ion and the interaction with ligand is strengthened by an aromatic residue on the receptor that forms van der Waals interactions with the aliphatic portion of the lysine residue that is docked in the “acidic pocket” (Fisher et al., 2006) and by other lysine residues on ligands that form weak electrostatic interactions (Dolmer et al., 2013). This model has been substantiated by other LDL receptor family members in complex with ligands (Guttman et al., 2010; Jensen et al., 2006; Lee et al., 2010; Verdaguer et al., 2004; Yasui et al., 2007). Interestingly Rauch et al. (Rauch et al., 2020) found that chemical modification of lysine residues on tau prevented uptake of tau. An evaluation of the cryo-EM structure of paired helical filaments of tau isolated from AD brains (Fitzpatrick et al., 2017) revealed 6 surface accessible lysine residues (Willard et al., 2003) per monomer, which are displayed in Fig 7. Optimal distances between lysine residues for docking into CR repeats can be estimated from known structures of various CR repeats from this family of receptors, with a range of ~14-41 Å and average of 26 Å (Supplemental Table I). A number of surface accessible lysine residues on the filament core of tau isolated from AD patients are within this optimal distance for LRP1 binding, which supports the observation that LRP1 is capable of enhancing tau seeding.

**Figure 7.**
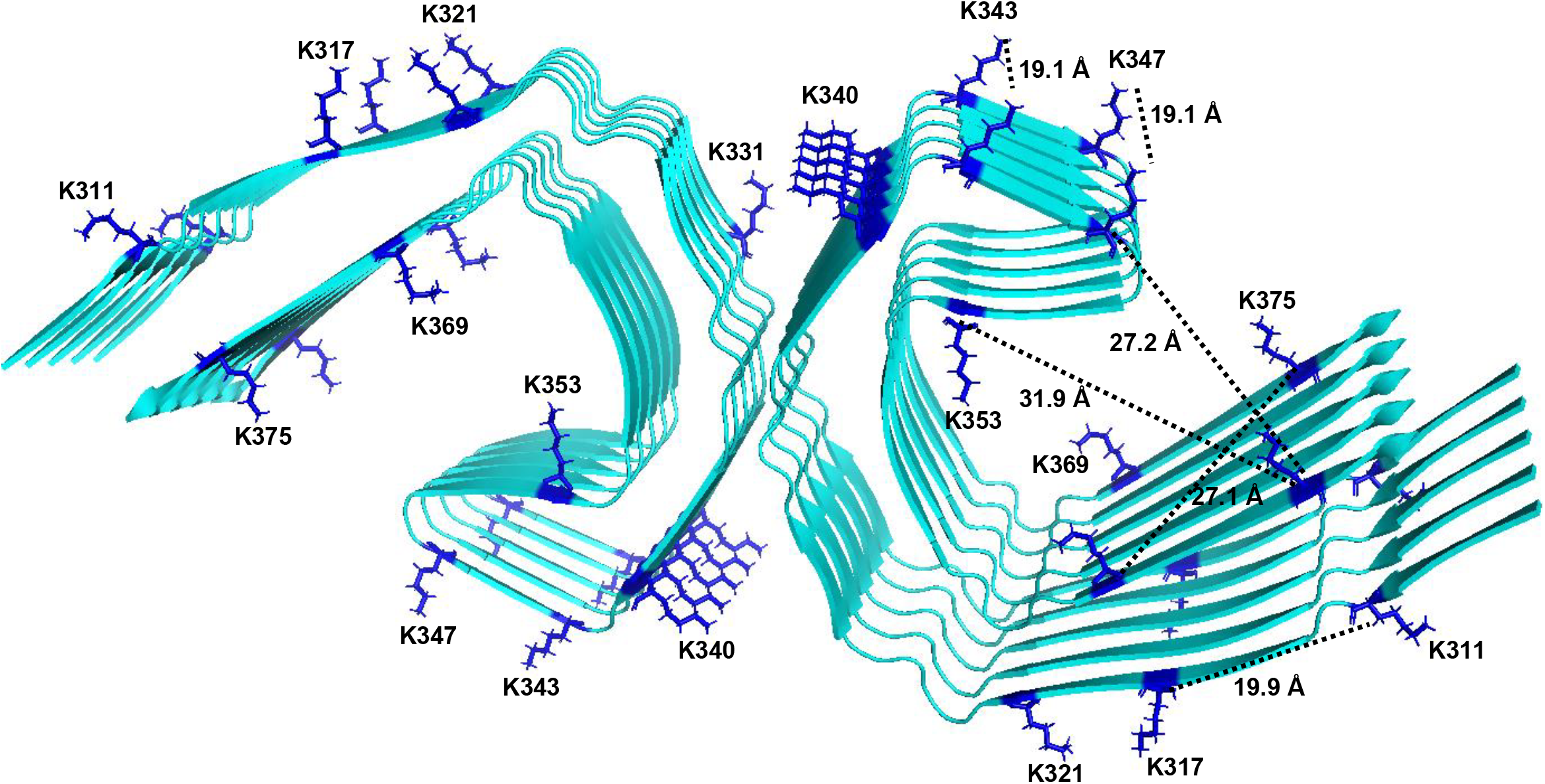
Surface accessible lysine residues available for LRP1 binding on tau protofilament from Alzheimer’s Disease. Ribbon diagram of tau filament core (PDB 5O3L)(Fitzpatrick et al., 2017) showing accessible surface areas for lysine residues available for interacting with LRP1 (ASA > 0.5). Accessible Surface Area was calculated from the coordinates in PDB 5O3L using VADAR(Willard et al., 2003). Distances between the a-carbon of lysine residues was determined using PyMOL software.

A recent study by Dujardin et al. (Dujardin et al., 2020) demonstrates that molecular diversity of tau contributes to AD heterogeneity and that some post-translational modification sites are associated with both enhanced seeding activity and worse clinical outcomes, while others are not. Our results demonstrate that phosphorylation of tau reduces its affinity to LRP1, suggesting LRP1 may differentially regulate various forms of tau, thus potentially providing a molecular basis for these differences. Changes in binding affinity may result in reduced internalization efficiency or altered intracellular trafficking of ligand in the endosomal compartments. Understanding the impact of various post-translational modifications on LRP1-mediated tau processing may be key to understanding the clinical and histopathological diversity of AD.

Our studies also determined that LRP1-independent mechanisms exist for the internalization of tau. We identified SORL1 as an additional receptor capable of binding and mediating the endocytosis of tau. Further, our data also reveal that expression of SORL1 in LRP1-deficient cells promotes tau seeding as well. SORL1 is a member of the VPS10P-domain containing receptor family that is involved in the trafficking of proteins among the Golgi apparatus, cell surface and endosomes. Genetic evidence suggests that variants of the *SORL1* gene are associated with AD (Holstege et al., 2017; Pottier et al., 2012; Rogaeva et al., 2007), and genetic deficiency of *SORL1* in mice results in increased Aβ levels (Andersen et al., 2005) and exacerbates early amyloid pathology (Dodson et al., 2008; Rohe et al., 2008) in mouse models of AD. Both LRP1 and SORL1 impact the trafficking of amyloid precursor protein (APP), but while the association of APP with LRP1 leads to enhanced amyloidogenic processing of APP and increases production of the Aβ (Pietrzik et al., 2002; Ulery et al., 2000), association of APP with SORL1 leads to a reduction in the amyloidogenic processing of APP (Andersen et al., 2005) by facilitating the trafficking of APP from endosomes to the Gogli apparatus (Schmidt et al., 2007). Thus, while a major function of LRP1 is to efficiently deliver cargo to lysosomal compartments, the major function of SORL1 is to deliver cargo between endosomes, the Golgi apparatus, and the cell surface. The specific mechanism(s) by which SORL1 contributes to tau seeding, like LRP1, will require further studies.

The N1358S mutation in the *SORL1* gene was identified in an exome sequencing study of patients with early onset AD (Pottier et al., 2012). Until now the functional consequences of this mutation have not been identified, though it has been proposed that this mutation could impact APP amyloidogenic processing (Mehmedbasic et al., 2015). While the most straightforward hypothesis would be that AD-associated genetic changes in SORL1 would enhance tau uptake and seeding, in fact our current analysis measuring recombinant tau uptake and AD-derived tau seeding in cell-based assays do not support this simple interpretation, as we find that the N1358S mutation results in impaired SORL1-mediated tau uptake and seeding. While we don’t fully understand the reason why we see reduced tau uptake and seeding with mutant SORL1, we nonetheless find it striking that SORL1 and apoE, two major AD genetic risk factors, are both implicated in tau uptake and seeding (Caglayan et al., 2014; Yajima et al., 2015). We speculate that there may be alternative pathways of pathological tau uptake in the brain, or that intracellular trafficking of tau, once taken up into the endolysosomal pathway, mediated by either LRP or SORL1, may be influenced in neurons by mechanisms that we are not modeling in CHO and HEK cells in culture; these pathways will be important to explore in future experiments.

As a final note, HEK293T cells stably expressing the FRET biosensor (Holmes et al., 2014) for tau aggregation have become the standard for assessing seed-competent tau, but require a protein transduction agent such as lipofectamine to efficiently demonstrate tau seeding. Our data demonstrate that HEK293T cells express low levels of LRP1 and do not effectively internalize tau or seed tau aggregation, but transfection of these cells with human LRP1 or SORL1 restores the ability of HEK293T cells to seed tau (even without lipofectamine) when incubated with pathogenic forms of tau.

In summary, our results demonstrate that LRP1 and SORL1 are both central receptors that regulate trafficking and metabolism of several important molecules linked to AD which include APP (Kounnas, Moir, et al., 1995; Ulery et al., 2000; Waldron et al., 2008) and β-amyloid (Shibata et al., 2000; Storck et al., 2016), tau, and apoE (Beisiegel et al., 1989; Kowal et al., 1989). Our work positions these two molecules as an unprecedented molecular point of convergence for the pathological hallmarks of AD. Furthermore, our findings show that they mediate both tau degradation and tau seeding, suggesting the role of LRP1 in tau pathology is nuanced and likely varies depending on the cellular environment and the species of tau involved. Understanding the individual pathways of each molecule and how they interconnect to LRP1 and SORL1 is key to the development of potential therapeutic intervention in Alzheimer’s disease and potentially other tauopathies.

## MATERIALS AND METHODS

### Cells

Human primary fibroblasts (WI-38) and human embryonic kidney (HEK293T) cells were purchased from ATCC and maintained in Dulbecco’s modification of Eagle’s medium (DMEM; Corning 10-013-CV) supplemented with 10% fetal bovine serum (FBS; Sigma F-4135). Wild-type Chinese hamster ovary (WT CHO) and CHO 13-5-1 cells (FitzGerald et al., 1995) were maintained in DMEM/Ham’s F12 with L-glutamine (DMEM/F12; Corning 10-090-CM) supplemented with 10% FBS. CHO cells deficient in xylosyltransferase (CHO-745) (Esko et al., 1985) provided by Jeffrey Esko (San Diego) were maintained in Kaighn’s Modification of Ham’s F-12 Medium (ATCC 30-2004) supplemented with 10% FBS. SH-SY5Y neuroblastoma cells were maintained in DMEM/F12 supplemented with 10% FBS. H4 neuoroglioma cells were maintained in DMEM supplemented with 10% FBS. Mouse embryonic fibroblasts (MEF) and PEA-13 cells were maintained in DMEM/F12 supplemented with 10% FBS. The Tau RD P301S FRET Biosensor embryonic kidney 293T cells (ATCC CRL-3275) provided by Marc Diamond were maintained in DMEM supplemented with 10% FBS. All cells were cultured with 1X penicillin-streptomycin (P/S; Corning 30-002-CI), and maintained at 37°C, 5% CO2 in a humidified atmosphere.

### Proteins, antibodies, and plasmids

Full-length human LRP1 was purified from placenta (Ashcom et al., 1990). Receptor associated protein (RAP) was expressed in *E. coli* (Williams et al., 1992). The mouse anti-LRP1 monoclonal antibody 5A6 was used to recognize the 85 kDa light chain of LRP1, and the rabbit anti-LRP1 polyclonal (R2629) antibody was used to inhibit ligand binding to LRP1 as previously described (Strickland et al., 1990). Full-length tau (2N4R; SP-495) and tau microtubule binding domain (MBD; SP-496) were purchased from R&D Systems. Recombinant human LRP1 cluster II, III, and IV Fc chimera proteins were produced by Molecular Innovations. His-tagged recombinant human tau variants 2N4R, 2N3R, and mutated proteins were expressed in *E. coli* or SF9 insect cells and purified. His-tag was removed by enzymatic cleavage. 2N4R tau harboring the 6A or 6E mutations were generated by converting T181, S199, S202, S396, S400 and S404 to alanine (6A mutant) or glutamic acid (6E mutant). Alpha-2-macroglobulin was purchased from Athens Research and was activated by methylamine as described (Ashcom et al., 1990). Full-length human *LRP1* was cloned into pN1 expression vector by VectorBuilder. Full-length human *SORL1* in pcDNA3.1 expression vector was provided by Claus Petersen(Jacobsen et al., 2001) and the N1358S mutant was generated by VectorBuilder. Recombinant human SORL1 aa 82-753 protein was purchased from R&D Systems.

### Tau internalization and degradation assay

Cellular internalization assays were conducted as previously described (FitzGerald et al., 1995; Kounnas, Moir, et al., 1995; Ulery et al., 2000). Twelve-well culture dishes were seeded with WI-38 (0.5×10^4^ cells per well), CHO (2×10^5^ cells per well), H4 (2×10^5^ cells per well), or HEK293T (9.5×10^4^ cells per well) cells. Cells were cultured overnight in DMEM (WI-38 and H4) or DMEM/F12 (CHO) with 10% FBS and 1X P/S. The following day, cells were incubated in assay media (DMEM supplemented with 1.5% bovine serum albumin (BSA) and 20 mM HEPES) for 1 hour, and then incubated with assay media containing 20 nM ^125^I-labeled tau (2N4R, R&D Systems, Inc. SP-495) in the presence or absence of 1 μM RAP for specified times. In some experiments ^125^I-labeled tau was co-incubated with 100 μM chloroquine (Sigma C6628), 20 μg/mL heparin (Sigma H-3125), or 300 μg/mL R2629. For experiments assessing a single cycle of tau uptake, MEF or PEA-13 cells were plated at 2×10^5^ cells/well and incubated overnight as described above. Cells were then incubated for 1 hour at 4 °C in assay media containing 20 nM tau with or without 1 μM RAP. After incubation, cells were washed with DPBS and fresh assay media maintained at 37 °C was added to the cells to trigger endocytosis. Cells were incubated at 37°C for designated times, and then collected assay cell-associated, internalized, and degraded tau. RAP-sensitive tau uptake was calculated by subtracting the RAP-inhibitable uptake from the total.

### Transfections

CHO WT, CHO 13-5-1, or HEK293T cells were plated at 9.5×10^4^ cells per well in a twelve-well culture dish in culture media without antibiotics. 24 hours after plating, cells were transfected with *LRP1* in pN1 expression vector, *SORL1* in pcDNA3.1 vector, or *SORL1* N1358S in pcDNA3.1 vector using 0.75μg DNA per well via PEI transfection reagent at a ratio of 6 μL PEI: 1 μg DNA. Transfection with empty vector was used as control. Cells were incubated overnight in transfection reagent, and 24 hours post-transfection the tau internalization assay was performed as described above.

### Surface plasmon resonance (SPR)

Binding of tau isoforms 2N4R, 2N3R, 2N4R tau harboring the 6A and 6E mutations, and hyperphosphorylated 2N4R tau produced in Sf9 cells to LRP1 or the VSP10 domain of SORL1 were assessed using a Biacore 3000 optical biosensor system (GE healthcare Life Sciences) essentially as described (M. Migliorini et al., 2020). Single kinetic titrations were performed by serial injections from low to high concentration (3.8, 11.5, 34.4, 103.3, 310 nM) with a 3.5 min injection time. Between sample runs, sensor chip surfaces were regenerated with 15 second injections of 0.5% SDS at a flow rate of 100 μl/min. In some experiments, sensor chip surfaces were regenerated with 15 second injections of 0.5% SDS at a flow rate of 100 μL/min with a low pH solution of 100 mM phosphoric acid (pH ~2.5).

### Enzyme-linked immunosorbent assays (ELISA)

ELISA were performed as previously described (Ulery et al., 2000). Briefly, microtiter wells were coated with 4 μg/mL human LRP1 in TBS overnight at 4°C, blocked with 3% BSA in TBS, then incubated with various concentrations (0 to 100 nM) of human recombinant tau (2N4R, R&D Systems Inc.) in the absence or presence of RAP (at a 20-fold molar excess compared to the concentration of tau) in assay buffer (3% BSA, TBS 5mM CaCl_2_, 0.05% Tween-20) for 18 hours at 4°C. Tau binding was detected with mouse monoclonal anti-tau IgG (Santa Cruz SC-21796).

### Tau seeding FRET biosensor assay

Human brain homogenates were prepared from an AD Braak VI brain and one healthy control brain from the Massachusetts Alzheimer’s Disease Research Center Brain Bank. Briefly, 100mg of frontal cortex tissue (Brodmann area 8/9) were thawed and homogenized in 500μl of PBS with protease inhibitor (Roche) by 30 up and down strokes in a glass Dounce homogenizer. The homogenate was centrifuged at 10,000 g for 10 min at 4°C. The supernatant was aliquoted and a bicinchoninic acid assay (BCA, Thermo Scientific Pierce) was performed according to manufacturer’s instructions to quantify total protein concentration. Soluble high-molecular weight seeding-competent tau (HMW SEC tau) was isolated from homogenate using size exclusion chromatography on a Superdex200 10/300GL column (#17-5175-01, GE Healthcare) as described previously(Takeda et al., 2015). Total tau concentration was measured by ELISA (# K15121D, Meso Scale Discovery). The seeding assay that had been previously described was adapted for the present study (Furman et al., 2015; Holmes et al., 2014). CHO WT and CHO 13-5-1 cells were reverse transfected with a pcDNA3 plasmid containing a construct that encoded the 344-378 residues of human P301L mutant tau fused to mTurquoise2, a self-cleaving 2A peptide, and 344-378 of human P301L mutant tau fused to Neon Green in Costar Black (Corning) clear bottom 96-well plates, using trans-IT X2 reagent (Mirus) according to manufacturer’s protocol. Cells were seeded at 60,000 cells/well and transduced with 100ng DNA/well. The next day, transfection media was replaced with 50μl of OptiMEM containing 50ng of total tau from human brain extract in the presence or absence of RAP. For positive controls, brain extracts were incubated with 1% lipofectamine 2000 for 15 min and then added to the wells, forcing the entry of tau seeds. Each condition was tested at least in quadruplicate. Cells were incubated with lysates for 24 to 28 hours. Cells were then collected using trypsin and transferred into 96-well U-bottom plates (Corning) using 10% FBS culture media to neutralize trypsin. Cells were pelleted at 1200 g for 10 minutes, resuspended in cold 2% paraformaldehyde for 10 minutes, pelleted at 1200 g and resuspended in 200μl of PBS. Samples were run on the MACSQuant VYB (Miltenyi) flow cytometer for the quantification of Turquoise fluorescence and Forster resonance energy transfer (FRET). Tau seeding was quantified by multiplying the percent of FRET-positive cells by the median fluorescence intensity of those cells, as described previously (DeVos et al., 2018). 40,000 cells per well were analyzed. Data was analyzed using FlowJo software. For experiments in the Tau RD P301S FRET Biosensor embryonic kidney 293T cells, cells were plated at 30,000 cells/well in 1/20 poly-D-lysine (Merck Milipore) precoated Costar Black (Corning) clear bottom 96-well plates. After 24 hours, cells were transfected with 100ng/well of LRP1 in pN1 expression vector or SORL1 in pcDNA3.1 vector with 0.4ul/well lipofectamine 2000 (Themo Fisher). The following steps were similar to the assay in CHO cells. All competing reagents and drugs including RAP (1μM), R2629 anti-LRP1 (300μg/ml), and chloroquine (100uM, Sigma-Aldrich) were incubated on cells with brain homogenate.

### SDS-Page and Western Blot

Cell cultures were collected in RIPA lysis buffer and analyzed by western blotting as previously described. Equal amounts of protein from each sample was mixed with loading buffer with or without 100mM/L dithiothreitol, boiled for 5 minutes, resolved by electrophoresis on a NovexTM 4-12% Tris-Glycine Mini Protein Gel, and transferred to polyvinylidene difluoride membranes for western blot analysis. Membranes were blocked with Odyssey blocking buffer and incubated with anti-LRP1 (R2629 or 5A6) or anti-SORL1 (BD Biosciences) at a concentration of 1:1000 overnight at 4 °C. The membrane was washed three times with 0.05% Tween20 in tris-buffered saline (TBST), and the antibody binding to membrane was detected with IRDye® 680RD or 800 anti-mouse or anti-rabbit IgG secondary antibody (LI-COR Biosciences) at a concentration of 1:10,000. The membrane was then washed three times with TBST and imaged using a LI-COR Odyssey Infrared Imaging System.

### Experimental Design and Statistical Analysis

All results are represented as mean ± SEM or SD, as indicated. Data were analyzed for significance using two-tailed Student’s *t*-test, one-way ANOVA, or two-way ANOVA, with Tukey or Sidak multiple comparisons posttests, as indicated. A *P* value of < 0.05 was set as the threshold for significance.

## ACKNOWLEDGMENTS

This work was supported by grants R35 HL135743 (DKS) from the National Heart Lung and Blood Institute and from the Rainwater Trust (BTH), the JPB Foundation (BTH) and NIA R56AG061196 (BTH). JC, MW, and ALA were supported by T32 HL007698 while ALA was supported by F30 HL145952 from the National Heart Lung and Blood Institute. AL was supported by the Swiss National Science Foundation (P2ELP3_184403), the Alzheimer’s Association (AACSF-19-617308) and the Professor Dr Max Cloetta Foundation and Uniscientia Foundation, Vaduz.

## COMPETING INTERESTS

None

## AUTHOR CONTRIBUTIONS

JC, AL, DS, and BH conceived the idea and designed the study. JC designed, conducted, and analyzed uptake, SPR, ELISA, and immunofluorescence experiments. MM conducted and analyzed SPR experiments. AL designed, conducted, and analyzed tau seeding experiments. AA conducted immunofluorescence experiments and provided input to the manuscript. MW conducted alpha-2-macroglobulin uptake experiments and provided input to the manuscript. SD and SM provided input to experimental design and edited the paper. DS wrote the paper with input from all authors. SM contributed to this article as an employee of the University of Maryland, Baltimore. The content is solely the responsibility of the authors and do not necessarily represent the views of the National Institutes of Health or the United States Government.

---

**Supplemental Table I.**
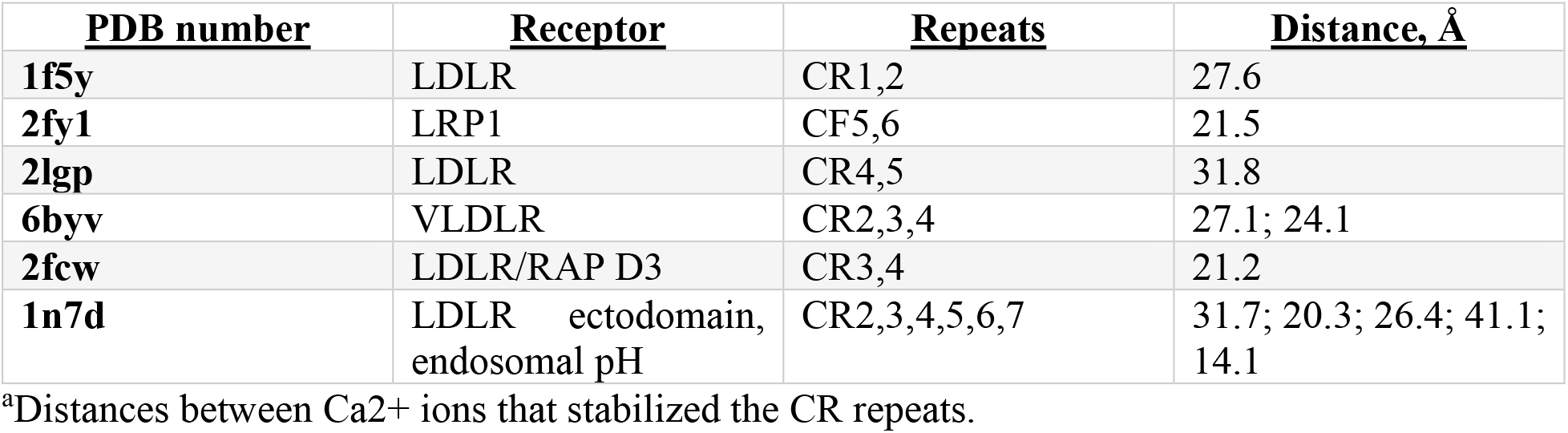
^a^Distances between complement-like repeats (CR) on LDL receptor family members

**Supplemental Figure 1.**
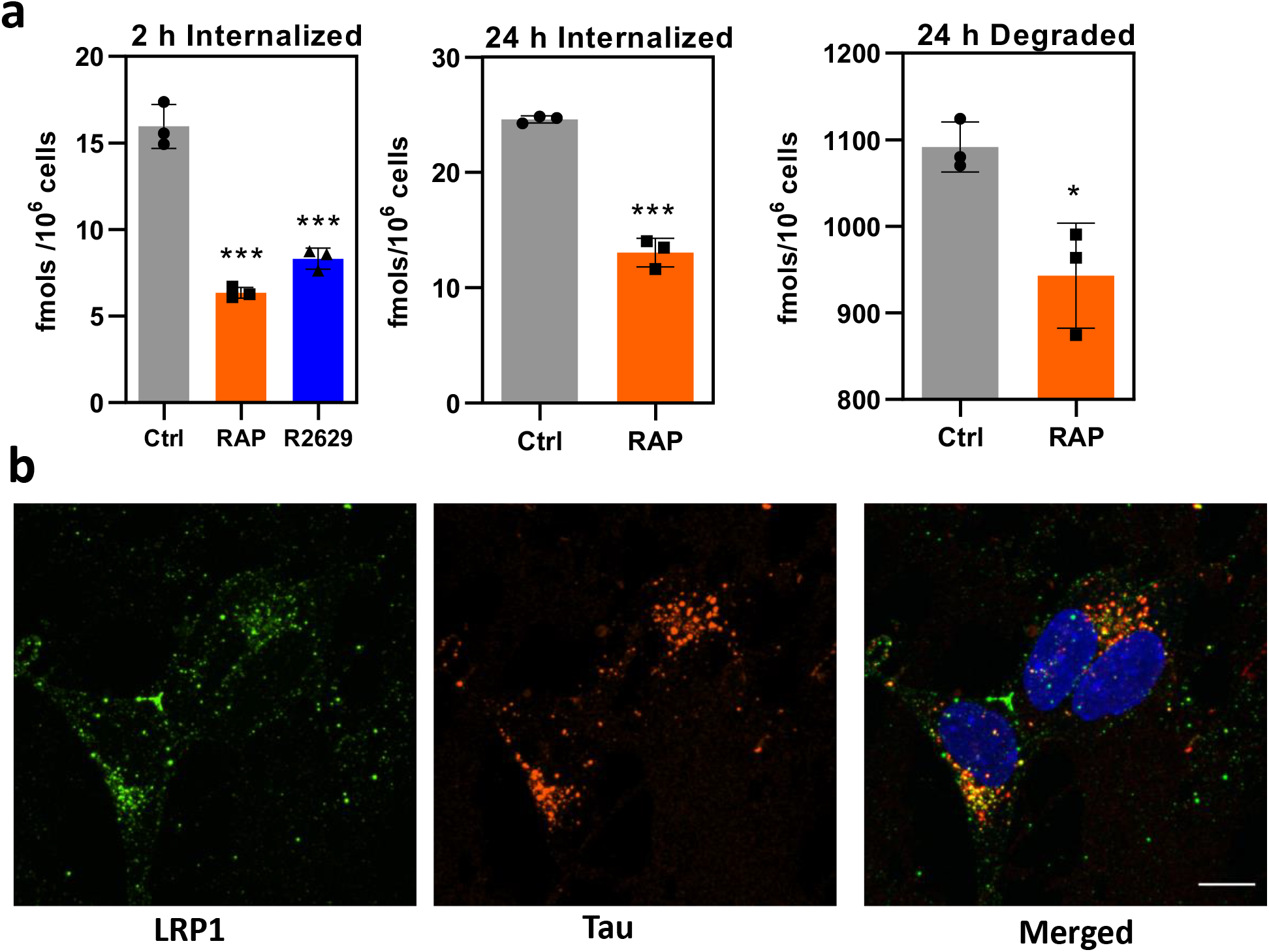
LRP1 mediates tau internalization and functional LRP1 colocalizes with tau in human neuronal cell lines. **(a)** H4 neuroglioma cells were incubated with ^125^I-labeled tau in the absence or presence of 1 μM RAP or 300 μg/mL anti-LRP1 IgG (R2629) for 2 or 24 hours at 37°C and the amounts of internalized, and degraded ^125^I-labeled tau were quantified. Data are expressed as mean ± SEM from three independent replicates. Statistical analysis was performed using one-way ANOVA (2 hr internalized: *F*(2,6)=113.2, *P* = 0.0001, followed by Sidak multiple comparisons (\*\*\**P*<0.0001) or t-test (24 hr internalized *n*=3, *P*=0.0001; 24 hr degraded (*n*=3, *P*=0.019). **(b)** Human neuroblastoma cells (SH-SY5Y) cells were grown on 8-chamber microscope slides until sub-confluent. The cells were serum starved by incubating with DMEM/F12 for 1 hour prior to experiment. The cells were then incubated at 37°C for 2 hours with monoclonal antibody 5A6 conjugated with AlexaFlour 488® (*green*) to label the endocytic pool of LRP1. After the cells were washed to remove unbound antibody, and they were incubated with 20nM tau conjugated with AlexaFlour 594® (*red*) incubated at 37°C for 2 hours. Colocalization of functional LRP1 and tau is displayed on merged panel (*yellow*). The scale bar is 10 μm.

**Supplemental Figure 2.**
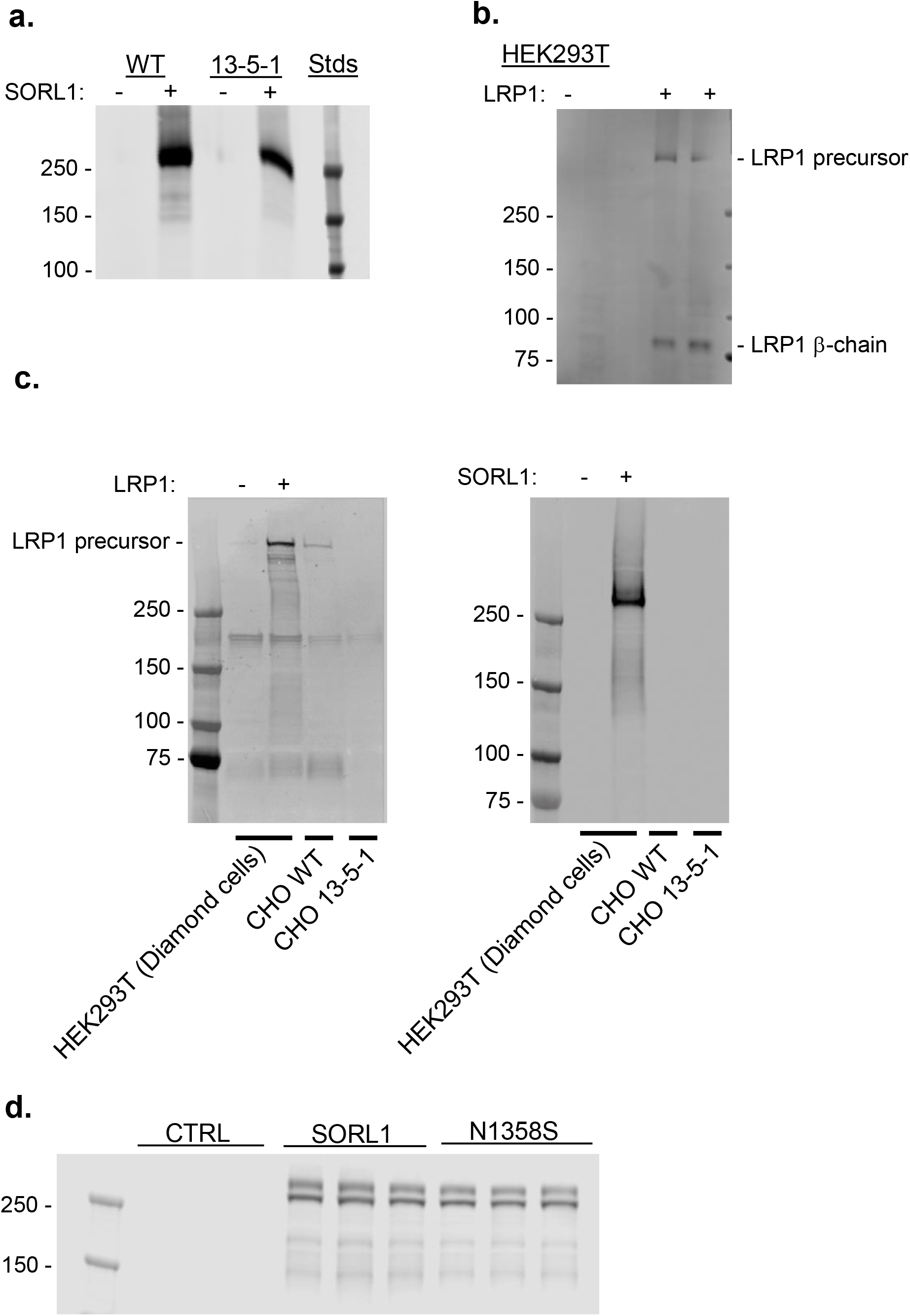
LRP1 and SORL1 expression in transfected cells. **(a**) CHO WT or 13-5-1 cells were transfected with SORL1 and collected 24 h post-transfection. Extracted proteins (non-reduced) were blotted for SORL1 using purified mouse anti-LR11 antibody (BD Biosciences Cat # 612633). **(b)** HEK293T cells were transfected with LRP1 and collected 24 h post-transfection. Extracted proteins were blotted for LRP1 using 5A6 anti-LRP1 antibody. **(c)** HEK293T FRET reporter Diamond cells were transfected with SORL1 and collected 24 h post-transfection. Extracted proteins were blotted for SORL1 using purified mouse anti-LR11 antibody (BD Biosciences Cat # 612633). **(d)** CHO 13-5-1 cells were transfected with SORL1 or N1358S SORL1 and collected 24 h post transfection. Extracted proteins (reduced) were blotted for SORL1 using purified anti-LR11 antibody.

## Supplemental data

### Immunofluorescence

SH-SY5Y cells were grown on 8-chamber microscope slides in DMEM/F12 supplemented with 10% FBS until sub-confluent. Cells were serum starved by incubating in DMEM/F12 in the absence of FBS for 1 hour prior to the experiment. 5A6 antibody and 2N4R tau were labeled using Alexa Flour® 488 and 594 antibody labeling kits according to manufacturer’s instructions (Life Technologies). LRP1 molecules undergoing endocytosis were labeled as previously described (Muratoglu et al., 2010) by incubating cells with 60 nM 5A6 anti-LRP1 antibody conjugated to Alexa Flour® 488 in DMEM/F12 at 37°C for 2 hours. Cells were washed to remove unbound antibody, and then incubated with 40 nM tau conjugated to Alexa Flour 594® in DMEM/F12 at 37°C for 2 hours. After washing to remove unbound tau, cells were fixed in 4% PFA for 10 minutes at room temperature. Slides were washed with DPBS and coverslips were mounted using VectaSheild Antifade with DAPI (Vector laboratories H-1200). Fluorescent images were acquired using a CSU-W1 spinning disk confocal system (Nikon, Yokogawa) in the Center for Innovative Biomedical Resources (CIBR) Confocal Microscopy Facility at the University of Maryland School of Medicine. Images were acquired with a 60x 1.49 NA oil-immersion objective as z-stacks with a step size of 0.1 μm and represented as maximal intensity projections along the z-axis in ImageJ software.

## Notes

### Competing Interest Statement

The authors have declared no competing interest.

